# Differential Short-Term Facilitation Of Synaptic Inputs And Spike Transmission At The Retinocollicular Synapse *In Vivo*

**DOI:** 10.1101/2024.01.17.576068

**Authors:** Kai Lun Teh, Elena Dossi, Nathalie Rouach, Jérémie Sibille, Jens Kremkow

## Abstract

Short-term plasticity (STP) is important for understanding how neuronal circuits can perform different computations. The STP of a neuron pair can be measured directly using paired whole-cell recordings. Besides, the cross-correlation between the presynaptic and postsynaptic neuronal firing is usually used as a proxy for estimating the synaptic properties. However, the relationships between the synaptic inputs and the spiking properties of the postsynaptic neurons during the STP *in vivo* still remain unclear. Here, we characterized the STP of both synaptic input, measured by the postsynaptic field potential (PFP), and spike transmission at the retinocollicular pathway of mice. We found that the STP of the retinocollicular pathway is mainly facilitating, where the second presynaptic spike induces a larger PFP and higher postsynaptic firing rate than the first presynaptic spike. The facilitation in the postsynaptic firing rate is generally larger than the PFP facilitation. Interestingly, the last postsynaptic spike timing also has a large facilitating effect on the postsynaptic spiking upon receiving a presynaptic input spike. However, the PFP does not depend on the last postsynaptic spike timing, suggesting that there is an input-independent component of spike transmission in STP. Overall, our results indicate that the STP of the retinocollicular pathway is likely a two-stage process, where the spiking plasticity of the postsynaptic neuron could be independent of its inputs.

**Highlights:**

- Measure the short-term plasticity of the postsynaptic dendritic response and the spike transmission simultaneously
- The retinocollicular pathway exhibits paired-spike facilitation
- Spike transmission facilitates more than postsynaptic dendritic response
- Short last postsynaptic spike time facilitates spike transmission independent of the next presynaptic input

## INTRODUCTION

The superior colliculus (SC) is a midbrain structure that is important for integrating multiple sensory modalities and driving behaviors, mainly visually guided behaviors (Ito and Feldheim, 2018; Isa et al., 2021; Basso et al., 2021). One important aspect in understanding the underlying circuit mechanisms of behaviors in the SC is short-term plasticity (STP), which likely underlies the functional dynamics of the SC circuits (Evans et al., 2018). STP have been studied for decades and is an important mechanism that plays a role in many brain functions, including temporal information processing of the neural circuits (Fortune and Rose, 2001), sensory adaptation (Chung et al., 2002), sound localization (Kuba et al., 2002; Cook et al., 2003), working memory (Mongillo et al., 2008), and escape decision (Evans et al., 2018).

In general, STP is difficult to be measured *in vivo* because it is challenging to target monosynaptically connected pairs *in vivo* (Jouhanneau et al., 2015). Nevertheless, STP has been characterized in different visual pathways *in vivo* using extracellular recordings, including the retinogeniculate (Usrey et al., 1998), thalamocortical (Stoelzel et al., 2008), and corticotectal (Bereshpolova et al., 2006) pathways. However, the STP in the retinocollicular pathway *in vivo* received less attention and remains poorly understood.

Besides, most STP studies focused either on the short-term dynamics of synaptic properties or spike transmission properties. There are only a few *in vitro* studies that examined the relationship between the synaptic inputs and the spiking outputs of the postsynaptic neurons during STP (Marder and Buonomano, 2003; Tominaga and Tominaga, 2016), where this relationship is largely unclear *in vivo*. We recently showed that retinocollicular pathway is a promising structure for simultaneous recording of the presynaptic axonal spiking, postsynaptic dendritic responses, and postsynaptic neuronal spiking between functionally connected neuron pairs *in vivo* using Neuropixels probe (Sibille et al., 2022a). We also discovered that the subsequent retinal ganglion cell (RGC) spike with a short interspike interval (ISI) can drive the SC spiking responses more efficiently (Gehr et al., 2023). However, the effects of the short RGC ISI on the postsynaptic dendritic response and the relationship between the postsynaptic dendritic response and the postsynaptic spiking are still unclear. Since inferring synaptic effects from extracellular spiking is challenging and not as straightforward as it sounds (Shein-Idelson et al., 2017; Ghanbari et al., 2020), an approach that can quantify the postsynaptic dendritic response and the postsynaptic spiking directly is needed to get an insight into their relationships.

Using the high-density electrode Neuropixels probe (Jun et al., 2017), here we show that it is possible to simultaneously measure the ongoing STP of both postsynaptic dendritic response, termed postsynaptic field potential (PFP), and spike transmission of the retinocollicular pathway. We characterized the STP in the retinocollicular pathway and demonstrated that the STP in this pathway is mainly facilitating, both in the postsynaptic dendritic inputs and postsynaptic neuronal spiking. Overall, this study shed a new light on the STP of the retinocollicular pathway, which could have important implications on some SC driven behaviors.

## RESULTS

In our previous study, we demonstrated the feasibility of recording the axonal-dendritic signals of the retinocollicular synapses and the postsynaptic SC neuronal spikes simultaneously (Sibille et al., 2022a). We now employed this approach to characterize the STP in the retinocollicular pathway.

In the experiment, a Neuropixels probe was tangentially inserted into the visual layers of SC to record the extracellular activities while visual stimuli were presented to the animals to induce visual responses in the retinocollicular pathway (Fig. 1A). With this approach, we can measure the axonal spikes of single RGCs and the corresponding PFP evoked in the SC dendrites by the RGC spike (Fig. 1B-C). More specifically, the PFP is an extracellular signal with net excitation elicited in dendrites of multiple postsynaptic SC neurons by the spike of a single presynaptic RGC axon (Sauve et al., 1995; Shein-Idelson et al., 2017; Sibille et al., 2022a).

**Figure 1:**
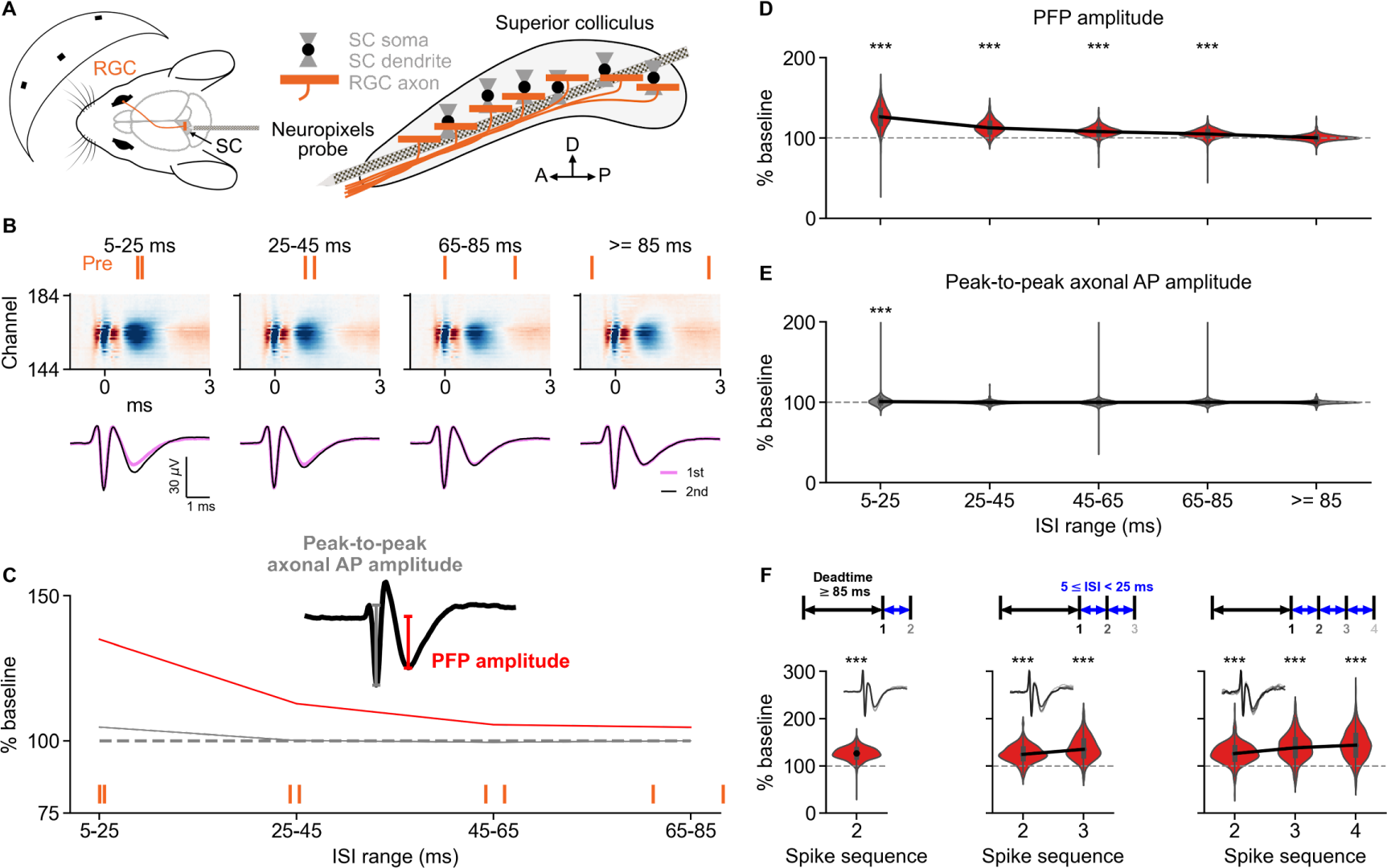
The PFP in SC facilitates reliably at short ISIs. (A) Scheme showing the experimental setup. Visual stimuli were presented to a mouse (left) with a Neuropixels probe tangentially inserted in the superficial layers of the contralateral SC (right). (B) Top: scheme showing different pre-pre ISI ranges. Middle: multichannel waveforms of an RGC correspond to the ISIs. Bottom: one-dimensional waveforms obtained by averaging across 11 channels (±5 channels from the best channel defined by Kilosort) of the multichannel waveforms. (C) The corresponding changes in PFP amplitude and peak-to-peak axonal AP amplitude between the first (baseline) and second RGC spikes for the waveforms in B. The PFP induced by the second RGC spike has a larger amplitude compared to the PFP induced by the first RGC spike for shorter ISIs while the peak-to-peak axonal AP amplitude remains largely unchanged for all ISIs. Inset indicates a typical recorded axonal waveform that has characteristic triphasic components (double negative peaks) with PFP amplitude and peak-to-peak axonal AP labeled. (D-E) The distributions of PFP amplitude (D) and peak-to-peak axonal AP amplitude (E) for different ISI ranges. The PFP amplitudes showed high facilitation in the ISI range of 5-25 ms, whereas the peak-to-peak axonal AP amplitudes are similar across different ISI ranges. The black line indicates the median, the box inside the violin plot represents the Q1 to Q3 of the data. (F) Two (same as D), three, and four consecutive RGC spikes within ISI of 5-25 ms induced lasting PFP facilitation. Top: scheme showing the different number of incoming presynaptic spikes after a dead time of at least 85 ms. Bottom: the percentage of the PFP amplitudes normalized to the PFP of the first spike. The black line indicates the median, the box inside the violin plot represents the Q1 to Q3 of the data.

### Shorter ISIs Induced Larger Postsynaptic Field Potential

To characterize the synaptic properties of the retinocollicular pathway, we analyzed the amplitude of the average PFP within five ranges of presynaptic-presynaptic (pre-pre) ISI, namely, 5-25, 25-45, 45-65, 65-85, and ≥85 ms. We found that the shorter the ISI, the larger the PFP amplitude evoked by the second RGC spike compared to the PFP amplitude evoked by the first RGC spike (Fig. 1D; Q1: 117.9%, 107.3%, 104.3%, 102.55%, 98.38%; median: 126.26%, 112.49%, 107.79%, 104.86%, 100.12%; Q3: 134.77%, 118.64%, 111.23%, 108.18%, 102.13%; *p* = 9.63 x 10^−79^, 3.34 x 10^−76^, 5.9 x 10^−68^, 4.32 x 10^−56^, 0.333 for ISI ranges of 5-25, 25-45, 45-65, 65-85, and ≥85 ms, two-sided Wilcoxon signed-rank test, *n* = 498 RGCs, *n* = 27 experiments, *n* = 24 mice). Similar effects were observed in the SC slice recordings (Fig. S1C).

As a control, the peak-to-peak axonal action potential (AP) amplitude remained largely unchanged between the first and second RGC spike (Fig. 1E; Q1: 99.25%, 99.01%, 99.15%, 99.25%, 99.2%; median: 100.85%, 99.88%, 99.95%, 100.03%, 99.93%; Q3: 102.68%, 100.7%, 100.72%, 100.76%, 100.71%; *p* = 1.91 x 10^−11^, 0.059, 0.199, 0.667, 0.293 for ISI ranges of 5-25, 25-45, 45-65, 65-85, and ≥85 ms, two-sided Wilcoxon signed-rank test, *n* = 498 RGCs, *n* = 27 experiments, *n* = 24 mice). Although the change of the RGC axonal AP amplitude of the 5-25 ms ISI range is significant compared to the baseline, its effect size is very small (Cohen’s d = 0.00909) and thus negligible.

To make sure that there are no confounding effects on the PFP signals by the postsynaptic SC neuronal spikes, we compared different variables of the average RGC waveforms with and without tailgating postsynaptic SC spikes. The average RGC waveforms without tailgating SC spikes were computed by removing the RGC spikes that have any detectable SC spikes occurring within the lag window [-3, 6] ms before averaging. No obvious difference in the PFP facilitation was found between the RGC waveforms with and without tailgating SC spikes for all variables quantified (Fig. S2; PFP amplitude: *p* = 0.904, 0.954, 0.365, 0.617 for ISI ranges of 5-25, 25-45, 45-65, and 65-85 ms, two-sided Wilcoxon rank-sum test, *n* = 110 RGCs and *n* = 9 mice for waveforms with tailgating SC spikes, *n* = 37 RGCs and *n* = 5 mice for waveforms without tailgating SC spikes). Moreover, comparison between the control condition and muscimol application in SC also showed no obvious difference in the PFP facilitation for all waveform variables quantified (Fig. S3; PFP amplitude: *p* = 0.046, 0.258, 0.23, 0.401 for ISI ranges of 5-25, 25-45, 45-65, and 65-85 ms, two-sided Wilcoxon rank-sum test, *n* = 110 RGCs and *n* = 9 mice for control, *n* = 44 RGCs and *n* = 3 mice for muscimol application). The application of muscimol in the SC inhibited the SC neuronal spiking, leaving the PFP unaffected by the SC spike waveform, and thus supporting the observation that PFP amplitude facilitates at shorter ISIs (Fig. 1D).

Since RGCs make strong and specific functional connections onto SC neurons (Sibille et al., 2022a) of both excitatory and inhibitory types (Gehr et al., 2023), the RGC axonal boutons might be overactivated and thus giving a small PFP facilitation (Fig. 1D) due to vesicle depletion. If this is the case, further RGC spikes might lead to depression instead. To determine whether the RGC axonal boutons are overactivated, we analyzed the PFP amplitudes induced by multiple consecutive RGC spikes with regular ISI ranges of 5-25 ms (Fig. 1F, top). We found that the PFP amplitude keeps increasing for at least three (Fig. 1F, middle; Q1: 114.68%, 121.21%; median: 124.4%, 135.04%; Q3: 137.06%, 153.58%; *p* = 2.41 x 10^−71^, 3.61 x 10^−76^ for the second and third spikes, two-sided Wilcoxon signed-rank test, *n* = 484 RGCs, *n* = 27 experiments, *n* = 24 mice) and four consecutive spikes (Fig. 1F, right; Q1: 114.49%, 121.93%, 122.51%; median: 126.11%, 138.31%, 143.96%; Q3: 139.53%, 156.62%, 165.1%; *p* = 2.28 x 10^−49^, 8.72 x 10^−51^, 1.27 x 10^−52^ for the second, third, and fourth spikes, two-sided Wilcoxon signed-rank test, *n* = 343 RGCs, *n* = 25 experiments, *n* = 22 mice) with similar ISIs in comparison to the PFP induced by the first RGC spike, suggesting that the observed facilitation can continue over multiple consecutive spikes. Again, similar observations were found in the SC slice recordings (Fig. S1D). In short, the larger PFP amplitude at shorter ISIs, together with the increasing PFP amplitude in several consecutive spikes, indicate that the retinocollicular synapses mainly undergo a small but reliable short-term facilitation.

### Shorter presynaptic ISIs Induce Larger Spike Transmission Probability

We have shown that short pre-pre ISIs could induce facilitation at the PFP level, but does this PFP facilitation translate into the spiking output of the postsynaptic SC neurons? To investigate the effects of the pre-pre ISI on the spike transmission, we quantify the excess spikes of the baseline corrected cross-correlation of connected RGC-SC neuron pairs after grouping the RGC spikes into the five ranges of pre-pre ISI (Fig. 2B, D; see Methods). The RGC axonal and SC neuronal spikes were distinguished based on their distinct waveforms (Fig. 2A; Sibille et al., 2022a, Gehr et al., 2023). The spike train cross-correlogram (CCG; Fig. 2C) of RGC-SC neuron pairs was then used for detecting potential connections and quantifying the spike transmission (Stark and Abeles, 2009; English et al., 2017).

**Figure 2:**
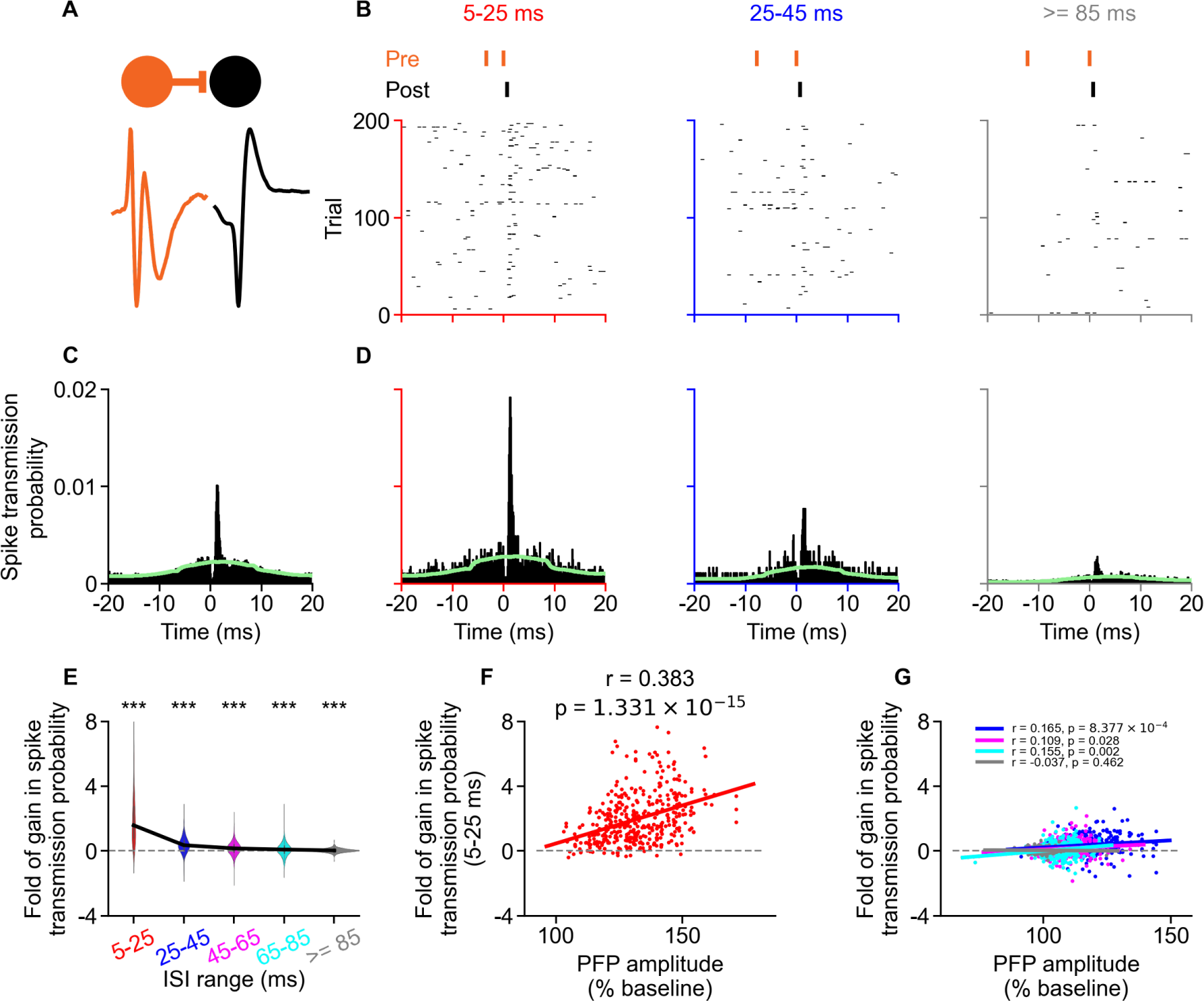
The PFP facilitation enhances the gain in spike transmission probability. (A) Scheme showing the presynaptic RGC (orange) and its postsynaptic SC partner (black) with their corresponding waveform. (B) Raster plots showing the postsynaptic SC activities upon receiving presynaptic inputs of different ISIs. (C-D) CCG showing the spike transmission probability of a connected pair averaged over all presynaptic RGC spikes (C) and averaged over RGC spikes of corresponding ISI ranges in B (D). The light green line indicates the CCG baseline used for computing the baseline corrected CCG. (E) The shortest ISI range (5-25 ms) has the largest fold of gain in spike transmission probability. The fold of gain in spike transmission decreases almost back to baseline at 45-65 ms. The black line indicates the median, the box inside the violin plot represents the Q1 to Q3 of the data. (F-G) The fold of gain in the spike transmission probability is positively correlated to the facilitation of the PFP amplitude. The ISI range of 5-25 ms has the strongest correlation (F) compared to other ISI ranges in G. ISI ranges of 25-45, 45-65, and 65-85 ms have small but significant positive correlation between the fold of gain in spike transmission probability and the PFP amplitude (G).

The gain in the spike transmission probability was computed by subtracting the spike transmission probability of the first RGC spikes from the second RGC spikes for each pre-pre ISI range. This gain was then normalized to the average pre-pre ISI spike transmission probability to obtain the fold of gain in the spike transmission probability. Similar to the PFP amplitude, the fold of gain in the spike transmission is also facilitating more at shorter pre-pre ISIs (Fig. 2E; Q1: 0.806, 0.079, ™0.074, ™0.123, ™0.033; median: 1.585, 0.356, 0.158, 0.088, 0.033; Q3: 2.747, 0.655, 0.392, 0.304, 0.099; *p* = 3.38 x 10^−67^, 6.36 x 10^−40^, 4.69 x 10^−19^, 5.34 x 10^−8^, 1.44 x 10^−8^ for ISI ranges of 5-25, 25-45, 45-65, 65-85, and ≥85 ms, two-sided Wilcoxon signed-rank test, *n* = 406 pairs, *n* = 224 RGCs, *n* = 21 experiments, *n* = 20 mice). The fold of gain in the spike transmission probability has a strong positive correlation to the facilitation of the PFP amplitude at 5-25 ms ISI range (Fig. 2F; *r* = 0.383, *p* = 1.33 x 10^−15^, *n* = 406 pairs), indicating that the facilitation of PFP amplitude can enhance the spike transmission probability for shorter ISIs. This suggests a presynaptic origin for the facilitation of postsynaptic spiking at 5-25 ms ISI range. Interestingly, the larger ISI ranges of 25-45, 45-65, and 65-85 ms also have a small but significant positive correlation between the fold of gain in the spike transmission probability and the STP of the PFP amplitude (Fig. 2G; *r* = 0.165, 0.109, 0.155, *p* = 8.38 x 10^−4^, 0.028, 0.002, respectively, *n* = 406 pairs). It is worth noting that the median PFP amplitude of the second RGC spike is around 126% of the first RGC spike for the 5-25 ms in Fig. 2F, which is a relatively small increase (∼0.26 fold increase) when compared to the median of ∼1.59 fold of gain in spike transmission probability. To sum up, spike transmission in the pre-pre ISI condition is facilitating and is strongly correlated to the PFP facilitation, although the large gain in spike transmission might not be entirely attributed to the small PFP facilitation.

### Shorter Post-Pre ISIs Facilitate Spike Transmission Probability Independent Of PFP

Next, we examined how the last postsynaptic spike timing influences the STP upon the arrival of a presynaptic spike (English et al., 2017). We did this to determine whether the activation of the postsynaptic SC neuron plays a role in converting the presynaptic input to spiking output. To answer this question, the interval from the last postsynaptic spike to the next presynaptic spike, given that the presynaptic spike has a dead time of at least 85 ms, was computed for each connected pair. This interval is termed postsynaptic-presynaptic (post-pre) ISI. Similar to the pre-pre ISIs, the post-pre ISIs were categorized into the five same ISI ranges.

In the post-pre ISI condition, there is a gap roughly equal to the smallest post-pre ISI of each group, which can be seen from the raster plots (Fig. 3A) and the corresponding CCGs (Fig. 3B). This ensures that for each ISI range, the connected postsynaptic neuron is silent for at least the smallest post-pre ISI. Our results show that the shorter the post-pre ISI, the higher the spike transmission probability upon the next presynaptic input (Fig. 3C; Q1: 0.466, 0.469, 0.46, 0.203; median: 1.408, 1.457, 1.053, 0.694; Q3: 3.108, 2.574, 2.205, 1.784; *p* = 5.12 x 10^−20^, 1.16 x 10^−20^, 8.05 x 10^−21^, 1.15 x 10^−19^ for ISI ranges of 5-25, 25-45, 45-65, and 65-85 ms, two-sided Wilcoxon signed-rank test, *n* = 131 pairs, *n* = 108 RGCs, *n* = 18 experiments, *n* = 17 mice).

**Figure 3:**
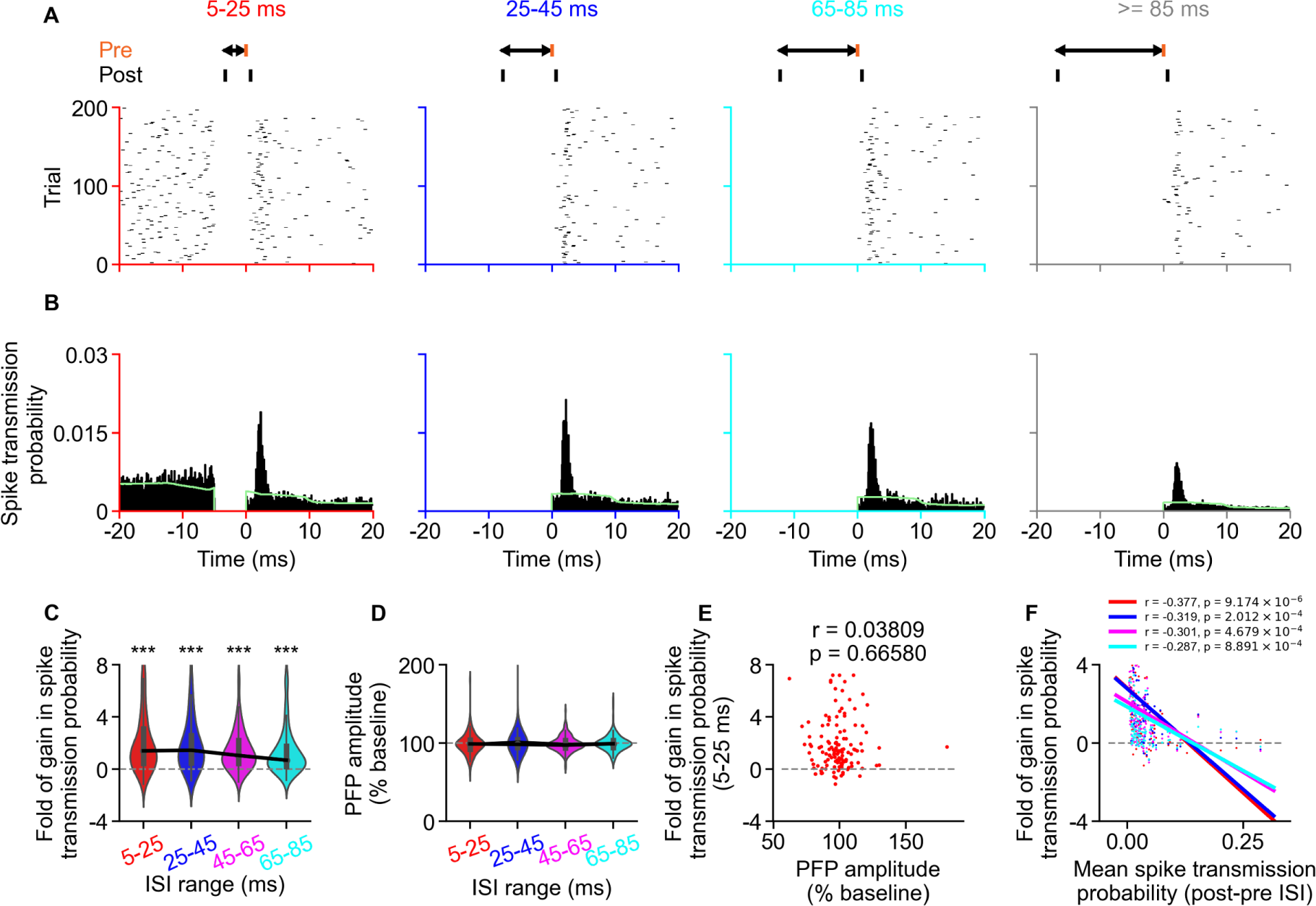
Postsynaptic spike timing induces spike transmission facilitation independent of the PFP amplitude. (A) Raster plots showing the postsynaptic activities upon receiving the presynaptic inputs of different post-pre ISIs. (B) CCGs showing the spike transmission probability of a connected pair computed with corresponding post-pre ISIs in A. The light green line indicates the CCG baseline. (C) The fold of gain in spike transmission probability normalized to the average post-pre ISI spike transmission probability against different post-pre ISI ranges. The black line indicates the median, the box inside the violin plot represents the Q1 to Q3 of the data. (D) No facilitation in the PFP for the post-pre ISIs. The black line indicates the median, the box inside the violin plot represents the Q1 to Q3 of the data. (E) There is no correlation between the PFP amplitude and the fold of gain in the spike transmission probability for the post-pre ISI condition. (F) The fold of gain in the spike transmission probability of the post-pre ISIs are negatively correlated to the average post-pre ISI spike transmission probability, with longer ISI ranges having less negative correlation.

Surprisingly, there is no facilitation observed in the PFP amplitude computed with the post-pre ISI condition (Fig. 3D; Q1: 92.2%, 94.32%, 94.2%, 94.56%; median: 99%, 98.47%, 97.58%, 99.2%; Q3: 103.86%, 104.84%, 103.39%, 103.77%; *p* = 0.178, 0.372, 0.078, 0.638 for ISI ranges of 5-25, 25-45, 45-65, and 65-85 ms, two-sided Wilcoxon signed-rank test, *n* = 131 pairs, *n* = 108 RGCs, *n* = 18 experiments, *n* = 17 mice), suggesting that the presynaptic input is likely not responsible for the increase in spike transmission under this condition. Indeed, there is no correlation between the fold of gain in the post-pre ISI spike transmission probability and the percentage facilitation of the PFP amplitude at 5-25 ms (Fig. 3E; *r* = 0.038, *p* = 0.666, *n* = 131 pairs). However, there are strong negative correlations between the fold of gain in the post-pre ISI spike transmission probability and the average post-pre ISI spike transmission probability for all ISI ranges (Fig. 3F; *r* = ™0.377, ™0.319, ™0.301, ™0.287, *p* = 9.17 x 10^−6^, 2.01 x 10^−4^, 4.68 x 10^−4^, 8.89 x 10^−4^ for ISI ranges of 5-25, 25-45, 45-65, and 65-85 ms, *n* = 131 pairs). This suggests that the RGC-SC neuron pairs that have a strong connection are less likely to facilitate in the post-pre ISI condition. In short, SC neurons have a higher spike transmission shortly after their previous spike, even though there is no apparent increase in the synaptic input, and this type of facilitation is more prominent in the weaker connections.

## DISCUSSION

Short-term plasticity (STP) has been studied extensively in many brain regions over the past four decades. However, STP in the retinocollicular pathway remains poorly understood. Since SC plays an important role in many behaviors essential for survival and an SC-related synaptic thresholding mechanism has been implicated in the defensive behaviors (Evans et al., 2018), characterizing the STP in this pathway could give some insights into the circuit dynamics of the SC that underlie those behaviors. Using the tangential Neuropixels probe recording in the mouse SC, a recently emerging model for the investigation of functional neural circuits, we demonstrate that the STP in the retinocollicular pathway is mainly facilitating, both in the postsynaptic dendritic response and the postsynaptic SC spiking.

### Simultaneous Recording Of PFP And Spike Transmission

Measuring STP *in vivo* by pairing the presynaptic and postsynaptic neurons has been a challenging task. One advantage of the *in vivo* tangential recording approach over the *in vitro* brain slice recording is that the STP can be measured more physiologically using visually evoked responses instead of electrically evoked responses. Besides, the neurons in an intact brain are in a more physiological environment than neurons in a slice. With waveform classification, we can distinguish the responses of individual neurons and axons (Sibille et al., 2022a). Furthermore, this technique has the potential to be applied in other brain regions, e.g., in the cortex (Sibille et al., in press). Last but not least, a large number of synaptic and spiking responses could be measured simultaneously due to the high-density electrodes of the Neuropixels probe.

### Facilitation In Spike Transmission Probability Is Much Larger Than In PFP Amplitude

Our results show that the facilitation of the spike transmission is much larger than the facilitation of the PFP. In the context of long-term potentiation (LTP), the higher facilitation in the population spikes compared to the facilitation in the field EPSP (fEPSP) has been observed more than three decades ago in the hippocampus (Bliss and Lomo, 1973; Taube and Schwartzkroin, 1988). Our results now suggest that this phenomenon also holds at the level of individual axons and individual postsynaptic neurons *in vivo*, where the PFP facilitation (Fig. 1D; median = 0.26 fold gain) is relatively small compared to the gain in the spike transmission (Fig. 2E; median = 1.59 fold gain). This suggests that, within a RGC-SC neuron pair, the gain in the postsynaptic spiking outputs evoked by the gain in synaptic inputs is nonlinear, illustrated by the decreasing gradient of the correlations between fold of gain in the spike transmission and the PFP plasticity (Figs. 2F-G).

### Input-Independent Facilitation In Spike Transmission

Spiking transmission has long been used as the proxy for inferring the synaptic properties (Ahissar et al., 1992; Constantinidis and Goldman-Rakic, 2002; Henze et al., 2002; English et al., 2017). However, to our surprise, the postsynaptic spiking activity is facilitated with no obvious increase in the presynaptic input during the post-pre ISI condition (Fig. 3E), suggesting other mechanisms might be at play during this condition. An explanation for this is the dynamic threshold (Azouz and Gray, 2000), where the spiking threshold of the postsynaptic SC neurons might change based on the past activity of that postsynaptic neuron. This suggests that there could be distinct STP effects for the synaptic and the spike transmission.

This phenomenon has been reported in the context of LTP, where there is a potentiation in the population spike amplitude without noticeable increase in the fEPSP (Bliss and Lomo 1973; Bliss and Gardner-Medwin, 1973). They also observed a decrease in the population spike latency, suggesting that the population spike potentiation is either due to a decrease in the spiking threshold (Chavez-Noriega et al., 1990) or increase in tonic activity from other pathways. With the advantage of measuring the spiking activity of individual postsynaptic neurons, we show that the facilitation in the spike transmission in our study is highly dependent on the interval to the last postsynaptic spike, indicating that a decrease in spiking threshold in the postsynaptic neuron is more likely to happen in our case than the tonic activity from other pathways.

This spike transmission facilitation in post-pre ISI condition is unlikely to be due to the inputs from other pathways, e.g. the corticotectal or the deeper-to-superficial SC connections, because those inputs need to be very precise temporally, strong, and have specific structure in order to contribute considerably to the time window used for detecting monosynaptic connections (Stevenson, 2023). However, we cannot rule out the possibility that the facilitation observed in the post-pre ISI condition could be due to the circuit-wide mechanisms, such as amplification within the SC (Shi et al., 2017), or brain states that modulate the gain of the SC neurons by changing the background synaptic inputs (Chance et al., 2002; Ferguson and Cardin, 2020). Another potential confounding effect that we did not take into account in this study is the converging RGC inputs into a postsynaptic SC neuron, which might contribute to the observed facilitation in the post-pre ISI condition, either directly to the postsynaptic neuron or indirectly via the intracollicular circuit amplification (Shi et al., 2017). However, since SC neurons are mainly driven by a strong RGC input and multiple weak RGC inputs (Sibille et al., 2022a), the facilitation in the post-pre ISI condition is less likely to be caused by converging RGC inputs.

### Relation To Previous Studies

STP has been characterized in the superficial SC of hamster SC slices (Balmer and Pallas, 2015), which was in the similar location as our experiments. However, by using the paired-pulse protocol, Balmer and Pallas (2015) observed short-term depression instead of facilitation, which contradicts our observations. To give a higher confidence to our observations, we performed similar experiments in mouse slices and observed that there is facilitation in paired-pulse protocol (Fig. S1C) as well as repetitive stimulation (Fig. S1D), similar to our *in vivo* results. The difference between the observations from Balmer and Pallas (2015) and ours is unclear, likely due to some cross-species differences.

### PFPs Are Unlikely To Be Contaminated By Postsynaptic Spikes

The waveforms of the PFP and the postsynaptic spikes can superimpose and thus bias the amplitude of the PFP (Sauve et al., 1995). To make sure the postsynaptic spikes do not contribute considerably to the PFP amplitude, we analyzed the plasticity of the PFP amplitude in two other conditions, namely 1) only analyze the PFP amplitude without postsynaptic spike within the time window from ™3 ms to 6 ms of the presynaptic RGC spikes (Fig. S2), and 2) analyze the PFP during the muscimol application that inhibited SC neuronal spiking (Fig. S3). In both conditions, the STP is similar to the normal condition without pharmacological application and with potential postsynaptic spikes. Another indirect evidence showing that the postsynaptic spikes are unlikely to contribute to the PFP is that there is no facilitation observed in the PFP of the post-pre ISI condition (Fig. 3D), even though there is a remarkable facilitation in the postsynaptic spiking activity (Fig. 3C).

### Limitations of the current study

Despite the fact that this approach allows us to measure the PFP and postsynaptic neuronal spiking simultaneously at a large scale, this approach also has some limitations. Although we could measure the PFP induced by a single axon, the PFP is the combined response of multiple postsynaptic dendrites that are connected to the axon (Sauve et al., 1995), and it is not yet known how to reliably disentangle the dendritic response of a single postsynaptic neuron. Also, the precise molecular mechanisms, both presynaptic and postsynaptic, that are responsible for the observed short-term facilitation cannot be determined without pharmacological applications. Lastly, we do not distinguish the RGC cell types (Baden et al., 2016) and SC cell types (Liu et al., 2023) in this study and it would be an important aspect to further refine our understanding of the STP on the cell-type specific level in future studies.

### Conclusions

Overall, our study shows that the STP at retinocollicular pathway is facilitating, both at the synaptic level and at the postsynaptic spiking level. However, the facilitation of the PFP cannot fully account for the facilitation of the spike transmission, consistent with previous findings in LTP. The tendency for a postsynaptic SC neuron to spike again shortly after the last spike suggests that mechanisms other than synaptic inputs might be at play, although the precise underlying mechanisms still need to be elucidated. Last but not least, our study demonstrates the feasibility to simultaneously record the STP at the synaptic and the somatic spiking levels *in vivo* using extracellular recordings, which opens the door for investigating the short-term circuit dynamics that is important for understanding the neural computations in different brain regions.

## METHODS

### Preparation, Animals, Surgery, And Histology

#### Animals

All experiments were conducted following the guidelines of the local authority (Landesamt für Gesundheit und Soziales - LAGeSo Berlin - G0142/18). Maximal care was provided to reduce the animal usage and their discomfort during all experiments. Adult male mice (C57BL/6J) were obtained from the local breeding facility (Charité-Forschungseinrichtung für Experimentelle Medizin, n = 20) and Charles-River Germany (n = 7).

#### Surgery

Induction was performed using 2.5% isoflurane in oxygen (Cp-Pharma G227L19A). Once stabilized, the animal was placed in a stereotactic frame (Narishige) with a closed-loop temperature controller (FHC-DC) for surgical procedure. During surgery, the eyes were protected with eye ointment (Vidisic) and the isoflurane level was slowly lowered (0.7-1.5%) while making sure that responses to tactile stimulation and vibrissa twitching are absent. A craniotomy was carried out shortly after constructing the dental cement-based crown (Paladur, Kuzler) used for fixing the head post and grounding. Once surgery and craniotomy were finished, the animal was transferred onto the recording table with a heating pad in place to keep the animal warm for the rest of the procedure. The experiment using visual stimuli (described below) was recorded once the Neuropixels probe is inserted tangentially and stabilized within the mouse SC.

#### Histology

Once the recording finished, the probe was removed, coated with DiI (Abcam-ab145311) diluted in ethanol, and re-inserted in the same tissue location for 5 minutes. The animal was then sacrificed with excess isoflurane (>4%). Phosphate buffer saline solution (PBS) was used for cardiac perfusions, followed by 4% paraformaldehyde (PFA) in PBS. The brains were kept in 4% PFA at least overnight and stored in PBS until histological slicing (Leica VT1200 S). DAPI-Fluoromount-G (70-100 µm slices, Biozol Cat. 0100-20) was used to mount the brain slices.

#### 3D location reconstruction

The Neuropixels probe track was reconstructed in 3D using SHARP-track (Shamash et al., 2018) and the Allen Mouse Brain Common Coordinate Framework (Steinmetz et al., 2019). The SC visually driven responses were used as landmarks to align and scale the estimated recording site positions.

### Visual Stimulation

#### Visual stimuli

All visual stimuli exposed to the animals were generated in Python using PsychoPy toolbox (Peirce, 2008). A TTL signal was generated and time-locked to the screen update for every stimulus onset. To cover a large part of the visual field, the visual stimuli were projected on a spherical dome (projector (NEC ME331W, refresh rate = 60 Hz, mean luminance = 110 cd/m², Gamma corrected) via a plexiglass reflecting half bowl (Modulor, 0260248). The inner part of the dome was covered with a layer of broad-spectrum reflecting paint (Twilight-labs) to increase the brightness of the reflected image (Denman et al., 2017). The projected image was warped using the meshmapper software (http://paulbourke.net/dome/meshmapper) to compensate for the deformation of the image reflections. In a subset of experiments (n = 4 mice), an LCD display (Dell S2716DG, refresh rate = 120 Hz, mean luminance = 120 cd/m^2^) was used instead of the spherical dome. The LCD display was positioned on the side of the animal and aligned to the pupil resting position in order to maximize the number of visually driven channels.

### Electrophysiological Recordings

#### Neuropixels recording

The neurons in the mouse SC were recorded with Neuropixels probes (Phase 3a and Phase 3B1) (Jun et al., 2017) using the Open Ephys software (www.open-ephys.org). All recordings were done with the PXIe system (National Instrument NI-PXIe-1071), which stores the extracellular signals in the local field potential (0.5-500 Hz) and the action potential bands (0.3-10 kHz).

#### SC tangential insertion

The Neuropixels probe was inserted using a micromanipulator system (NewScale, MPM-M3-LS3.4-15-XYZ Upright). Stereotactic coordinates were described in reference to lambda in the medio-lateral (ML), dorso-ventral (DV), and antero-posterior (AP) axes. All angles were defined in reference to the azimuthal plane at lambda (Paxinos and Franklin, Nixdorf 2007 stereotaxic atlas). The Neuropixels probe was inserted tangentially (15 to 25 deg) into the SC from the back (500 to 1200 μm ML, ™100 to ™500 μm DV, ™100 to ™300 μm AP from lambda) and remained parallel to the sagittal plane during the experiment. The Neuropixels probe was gradually driven >4 mm into the target tissue followed by a small withdrawal of 20 to 50 µm to release accumulated mechanical pressure. The probe was then left for ∼10-20 minutes to settle. To ensure that the probe is located within the area that is being visually stimulated, the receptive fields of the multi-unit-activity were mapped using a customized script (Sibille et al., 2022b) before any recording.

#### Waveform extraction and spike sorting

The extracellular neuronal signals were spike-sorted using Kilosort (2 and 2.5; Pachitariu et al., 2016). Manual curation using Phy2 (https://github.com/cortex-lab/phy) was performed on the Kilosort output. To avoid biases from noisy outputs, quality metrics thresholding, removal of unstable clusters, and controls for double-counted spikes were completed after the manual curation and before any further analysis. The identification of RGC axonal action potential waveforms was based on the characteristic presence of double negative peaks within 3 ms (Sibille et al., 2022a).

### Interspike Interval (ISI) Ranges

#### Pre-pre ISI

The pre-pre ISI is the ISI of two consecutive spikes of an RGC. For each RGC, if a pair of spikes has a dead time of at least 85 ms prior to the first spike, they will be categorized into one of the five ISI ranges: 5–25, 25–45, 45–65, 65–85, and ≥85 ms.

#### Post-pre ISI

The post-pre ISI of an RGC-SC neuron pair is the interval from the last occurring postsynaptic spike to the current presynaptic spike. To remove the potential effects of the pre-pre ISI, the post-pre ISI is valid only if there is a dead time of at least 85 ms before the presynaptic RGC spike time. The valid post-pre ISI was then categorized into one of the five ISI ranges: 5–25, 25–45, 45–65, 65–85, and ≥85 ms.

### Analysis of Postsynaptic Field Potential

#### PFP based on ISI range

The postsynaptic field potential (PFP) is the extracellular postsynaptic potential originated from the activation of multiple dendrites evoked by the spike of their same presynaptic RGC partner (Sauve et al., 1995; Sibille et al., 2022a). It is the second negative deflection following the RGC axonal spike. The one-dimensional PFP of an RGC was obtained by averaging 11 channels of the multi-channel waveform centered at the best channel defined by the Kilosort’s output (Fig. 1B). The PFP for each pre-pre ISI range was obtained from averaging the RGC spike waveforms within the same pre-pre ISI range. The PFPs evoked by the first and second spikes of the RGC spike pairs (with a valid dead time) within a pre-pre ISI range were averaged separately. Similarly, the PFP for each post-pre ISI range was obtained from the average RGC spike waveform of the post-pre ISI range. All RGCs were detected by Kilosort based on the first negative peak of the waveform, which corresponds to the extracellular potential evoked by the RGC axonal arbors in the SC (Sibille et al., 2022a). This ensured the detection of RGC spikes independent of any postsynaptic plastic effect.

#### PFP plasticity

Three variables were used for quantifying the PFP plasticity, namely PFP amplitude, peak-to-peak PFP amplitude, and PFP slope (Fig. S2A). The amplitude of the PFP was defined as the difference from the RGC axonal spike waveform baseline to the minimum point of the PFP (Fig. 1C). The peak-to-peak PFP amplitude was defined as the amplitude from the previous peak to the minimum point of the PFP and the PFP slope was defined as the slope from 0.2 to 0.8 fraction of the peak-to-peak PFP (Fig. S2A). The PFP of an RGC was included if (i) the number of RGC spike pairs for every ISI range is at least 500 (50 for the spike triplets and quadruplets in Fig. 1G-H, 200 for muscimol group in Fig. S3), (ii) its amplitude evoked by the second RGC spike at ≥ 85 ms pre-pre ISI is at least 5 μV in size, and (iii) the ratio of the PFP amplitude to the peak-to-peak PFP size is at least 0.3.

The plasticity in the PFP of a pre-pre ISI range, *T*, was defined as the percentage of PFP amplitude evoked by the second RGC spike, A^2*nd*,*T*^, over the PFP amplitude evoked by the first RGC spike, 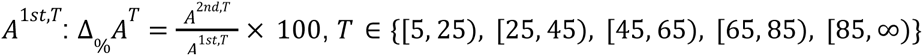 ms. For the post-pre ISI condition, the change in the PFP amplitude of an ISI range was defined as the percentage of PFP amplitude of a test ISI range, A^*T*^, over the PFP amplitude ≥85 ms, 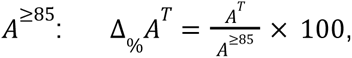 *T* ∈ {[5, 25), [25, 45), [45, 65), [65, 85)} ms.

### Cross-Correlation Analyses

#### Baseline corrected cross-correlation

Spike transmission was estimated from the spike train cross-correlogram (CCG) of an RGC-SC neuron pair. For unbiased estimation of the spike transmission, a slow co-modulated baseline, λ(*m*), was computed for each CCG, *C*(*m*), and subtracted from the CCG to get the baseline-corrected CCG, *Z*(*m*) = *C*(*m*) − λ(*m*) (Stark and Abeles, 2009; English et al., 2017). The baseline was computed by convolving a partially hollow Gaussian kernel (hollow fraction = 0.6, standard deviation = 10 ms, kernel length = 15 ms, bin width = 0.1 ms) with the CCG. The section at both ends of the CCG, with length equals to half the kernel length (7.5 ms), were duplicated, flipped, and symmetrically appended to the CCG when computing the CCG baseline to prevent any edge effects (Stark and Abeles, 2009). For the CCG of the post-pre ISI condition, there is a gap of the size of the smallest ISI, *Tmin*, within the post-pre ISI range. For example, there is a gap of 5 ms for the post-pre ISI range of 5-25 ms. This gap, with time window (− *Tmin*, 0] ms, was filled with the duplicated and flipped values from the post-pre ISI CCG window of [5, 5 + *Tmin*) ms when computing the post-pre ISI CCG baseline.

#### Connection detection

Connections are estimated from the excess count in the *m*th time lag of the CCG in comparison to the baseline using Poisson distribution with a continuity correction (Stark and Abeles, 2009; English et al., 2017). The RGC-SC pairs were labeled as connection if *P* < 0.001 for at least eight consecutive lag bins (bin width = 0.1 ms) within the lags of 0.8 - 2.8 ms.

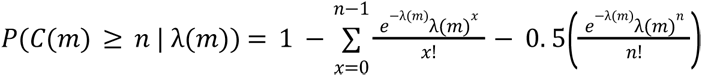

For analyses of the post-pre ISI condition, the post-pre ISI CCG was considered as having a significant effect if *P* < 0.01 for at least five consecutive lag bins.

### Spike transmission probability

#### Efficacy of synaptic connection

The spike transmission probability, *Pspike*, was computed from the rectified baseline-corrected CCG, (English et al., 2017) *Z* (*m*), averaged over *n* presynaptic spikes

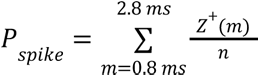

where *Z*^+^(*m*) = *max*(*Z*(*m*), 0)

#### Plasticity of spike transmission

The gain in spike transmission probability, *G*, was defined as the excessive spike transmission probability within lag window of 0. 8 − 2. 8 ms in a test group in comparison to the baseline group: 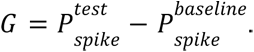. The fold of gain in spike transmission probability, *G_fold_*, was defined as the ratio of gain in spike transmission probability to the average spike transmission probability for the respective condition, 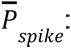 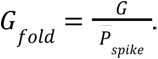

For the pre-pre ISI condition, the gain in spike transmission probability was defined as the excessive spike transmission probability in the second spike compared to the first spike of the RGC spike pair within an ISI range: 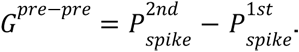 The fold of gain in *Pspike* for the pre-pre ISI condition, 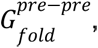, was defined as the ratio of *G*^*pre*−*pre*^ to the average spike transmission probability of the first and second spikes of all ISI ranges, 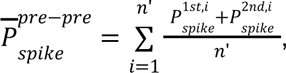, where *n*’ is the total number of valid spike pairs of an RGC. For the post-pre ISI condition, gain in *Pspike* was defined as the excessive spike transmission probability in an ISI range, *T*, compared to the baseline ISI range at ≥ 85 ms: 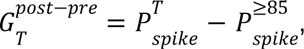 *T* ∈ {[5, 25), [25, 45), [45, 65), [65, 85)} ms. The fold of gain in *Pspike* for the post-pre ISI condition, 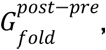, was defined as the ratio of 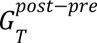 to the average spike transmission probability of all valid post-pre ISIs of an RGC-SC neuron pair.

### *In vitro* experiments and data analysis

#### Animal

Experiments were carried out according to the guidelines of the European Community Council Directives of 1 January 2013 (2010/63/EU). Mice (Mus musculus) were group housed on a 12-h light/dark cycle and all efforts were made to minimize the number of used animals and their suffering.

#### Slice preparation

Acute sagittal superior colliculus (SC) slices (400 µm) were prepared from 4-6 weeks-old C57BL/6 mice. Briefly, slices were cut at low speed (0.04 mm/s) and at a vibration frequency of 70 Hz in ice-cold oxygenated ACSF supplemented with sucrose (in mM: 87 NaCl, 2.5 KCl, 2.5 CaCl2, 7 MgCl, 1 NaH2PO4, 25 NaHCO3, and 10 glucose, saturated with 95% O2 and 5% CO2). Slices were maintained in a storage chamber containing standard ACSF (in mM: 119 NaCl, 2.5 KCl, 2.5 CaCl2, 1.3 MgSO4, 1 NaH2PO4, 26.2 NaHCO3, and 11 glucose, saturated with 95% O2 and 5% CO2) for at least 1 hour before recording.

#### Electrophysiological recordings

Slices were transferred in a submerged recording chamber mounted on an Olympus BX51WI microscope equipped for infrared differential interference microscopy and were perfused with standard ACSF at a rate of 2 ml/min at 34°C. Extracellular field potential (fEPSP) recordings were performed in SC superficial layers using glass pipettes (2-5 MΩ) filled with ACSF. Basal evoked postsynaptic responses were induced by stimulating SC afferents at 0.1 Hz. PPF was performed by delivery of two low stimuli (<10 mA) at various inter-pulse intervals (5-25, 30-50 and 60-100 ms). Repetitive stimulation was performed by delivery 2, 3 or 4 stimuli at 10 ms interval. Field potential recordings were acquired with Axopatch-1D amplifiers (Molecular Devices), digitized at 10 kHz, filtered at 2 kHz, and stored and analyzed on computer using pCLAMP 9 and Clampfit 10 software (Molecular Devices).

#### Statistics

Data are expressed as means ± SEM, unless otherwise stated. Repeated measures one-way ANOVA with Dunnett’s multiple comparison test was performed for PPF and repetitive stimulation comparisons.

## ACKNOWLEDGEMENTS

We thank Carolin Gehr and Tatiana Lupashina for helpful discussions during the project. We thank Francois David for comments on the manuscript. This work was supported by DFG Emmy-Noether grants KR 4062/4-1 and KR 4062/4-2 (J.K.).

## AUTHOR CONTRIBUTIONS

K.-L.T., J.S., and J.K. designed the study; J.S. conducted the *in vivo* experiments and curated the data; K.-L.T. analyzed the *in vivo* data; E.D. and N.R. designed and conducted the *in vitro* experiments; E.D. and N.R. analyzed the *in vitro* data; and K.-L.T. wrote the manuscript with inputs from J.S. and J.K.

## SUPPLEMENTARY MATERIALS

**Figure S1:**
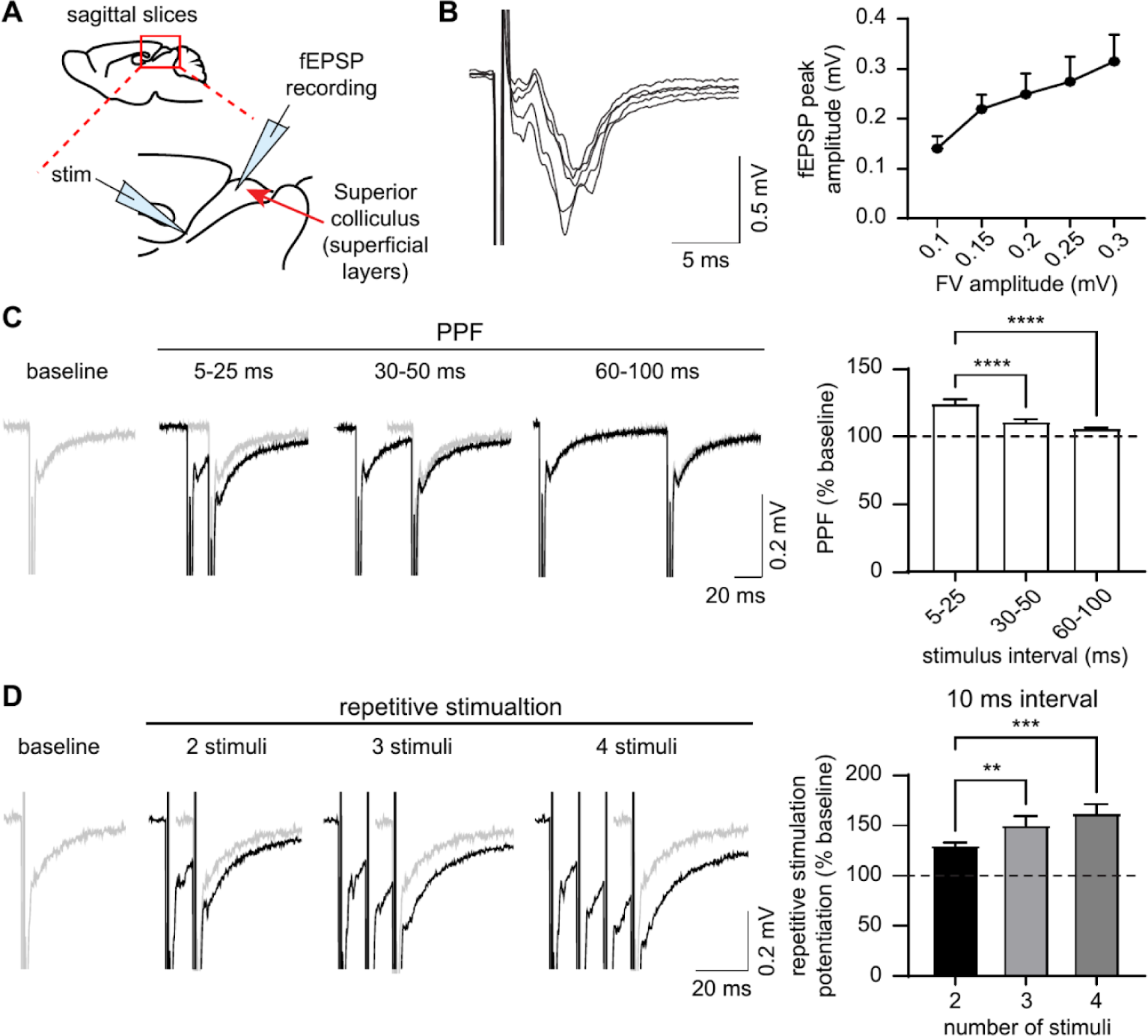
*In vitro* SC slices exhibit short-term facilitation in paired-pulse and repetitive stimulation protocols. (A) Schematic representation of electrode positions used in brain sagittal slices to record field excitatory postsynaptic potentials (fEPSPs) in the superficial layers of the superior colliculus (SC) evoked by electrical stimulation of SC afferents. (B) Input-output curves for basal synaptic transmission. Left: representative recordings in wild-type mice. Scale bars, 5 ms, 0.5 mV. Right: quantification of the fEPSP peak amplitude for different fiber volley amplitudes after stimulation (*n* = 7 slices from 3 mice). The data are represented as the mean ± sem. (C) Left, representative traces (black) of paired-pulse facilitation (PPF) with different inter-pulse intervals (5-25, 30-50 and 60-100 ms) recorded in the superficial layers of the SC after electrical stimulation of SC afferents. The baseline response to 0.1 Hz stimulation is shown in grey. Scale bars: 20 ms, 0.2 mV. Right, quantification of the PPF as % of the baseline response (*n* = 7 slices from 3 mice; *p* < 0.0001, RM-one way ANOVA). (D) Left, representative traces (black) of the response to repetitive stimulation (2, 3 and 4 stimuli at 10 ms interval) recorded in the superficial layers of the SC after electrical stimulation of SC afferents. The baseline response to 0.1 Hz stimulation is shown in grey. Scale bars: 20 ms, 0.2 mV. Right, quantification of the repetitive stimulation-induced potentiation as % of the baseline response (*n* = 6 slices from 3 mice; *p* = 0.007, RM-one way ANOVA). Asterisks indicate statistical significance (**, *p* < 0.01; ***, *p* < 0.001; ****, *p* < 0.0001).

**Figure S2:**
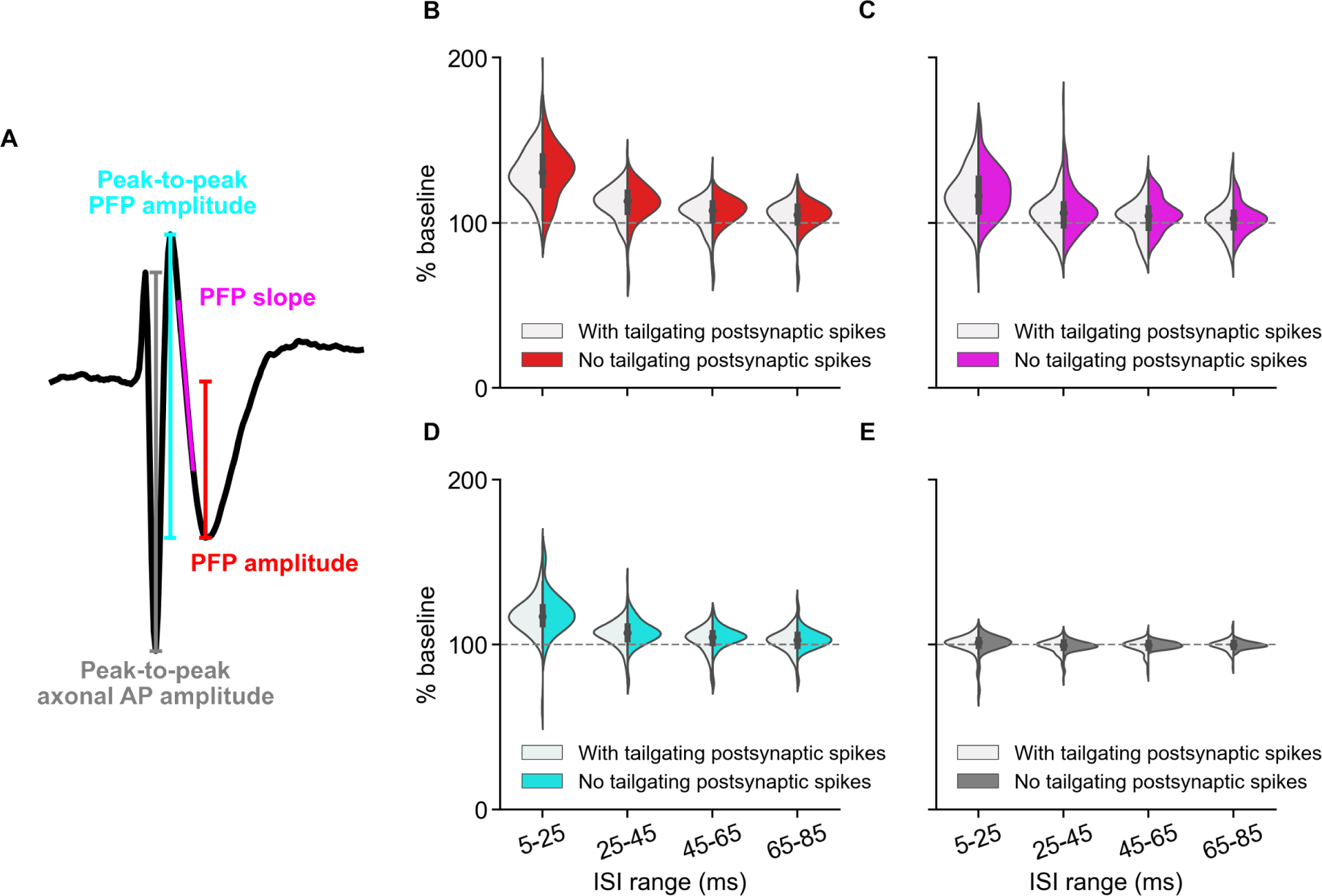
No significant difference in the PFP facilitation with and without tailgating postsynaptic SC spikes at the lag window [-3, 6] ms around the RGC spike time. (A) Scheme showing different variables of the 1-D RGC waveform being quantified. (B-E) The percentage change of different RGC waveform variables of the second RGC spike in different pre-pre ISI ranges in comparison to the baseline ISI range of ≥ 85 ms. No obvious difference observed between the waveforms with (white) and without (colored) tailgating postsynaptic SC spikes for facilitation in PFP amplitude (B; *p* = 0.904, 0.954, 0.365, 0.617), PFP slope (C; *p* = 0.363, 0.523, 0.251, 0.718), peak-to-peak PFP amplitude (D; *p* = 0.483, 0.872, 0.282, 0.602), and peak-to-peak axonal AP amplitude (E; *p* = 0.922, 0.668, 0.401, 0.738). For all variables, a two-sided Wilcoxon rank-sum test was used. For the group with tailgating postsynaptic spikes: *n* = 110 RGCs, *n* = 10 experiments, *n* = 9 mice. For the group without tailgating postsynaptic spikes: *n* = 37 RGCs, *n* = 5 experiments, *n* = 5 mice.

**Figure S3:**
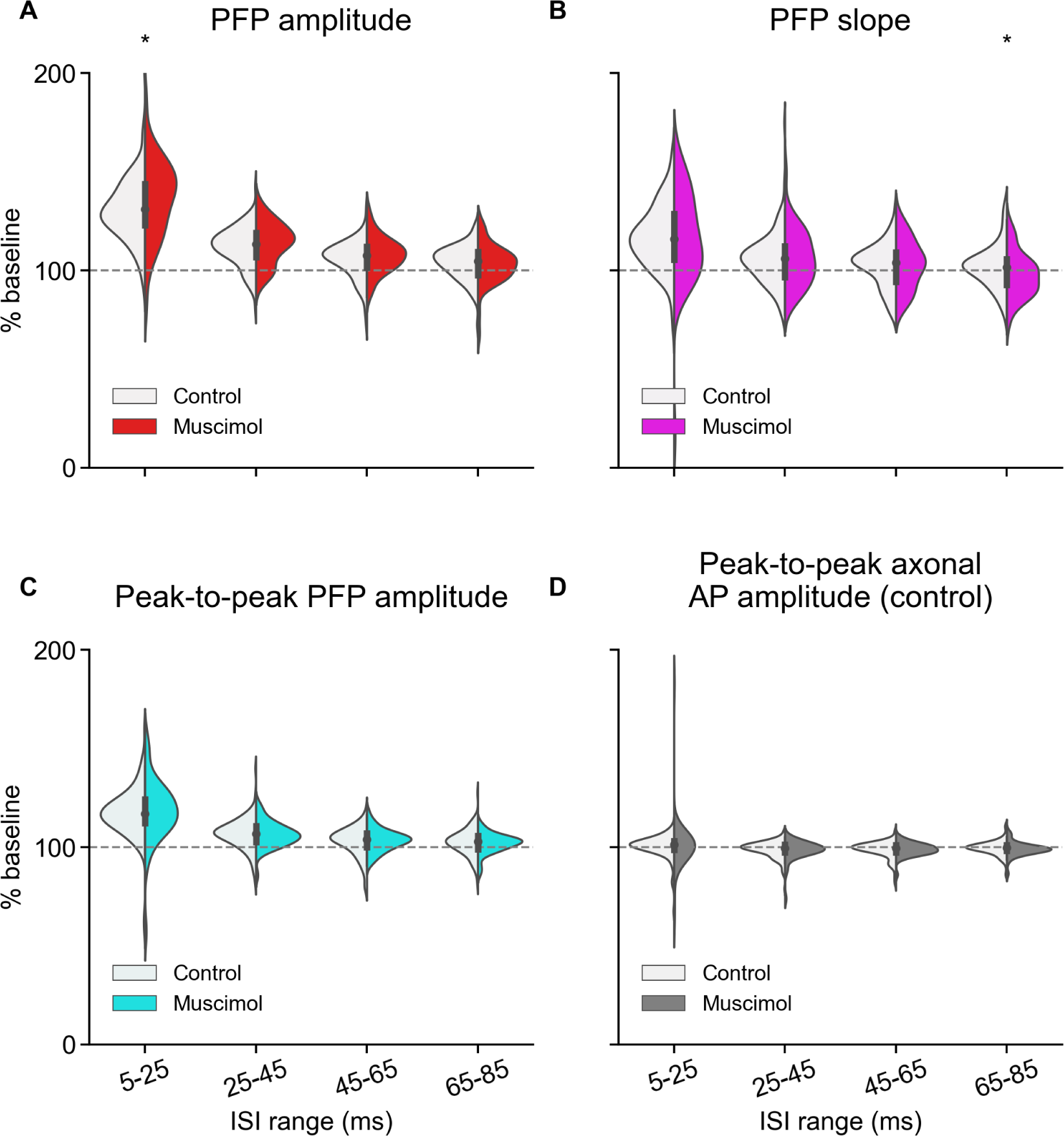
No obvious difference in the PFP facilitation between the control and with muscimol application in the SC. (A-D) The percentage change of different RGC waveform variables of the second RGC spike in different pre-pre ISI ranges in comparison to the baseline ISI range of ≥ 85 ms. No obvious difference observed between the controlled waveforms (white) and waveforms under muscimol condition (colored) for facilitation in PFP amplitude (A; *p* = 0.046, 0.258, 0.23, 0.401), PFP slope (B; *p* = 0.914, 0.901, 0.974, 0.016), peak-to-peak PFP amplitude (C; *p* = 0.541, 0.504, 0.974, 0.675), and peak-to-peak axonal AP amplitude (D; *p* = 0.614, 0.507, 0.598, 0.101). For all variables, a two-sided Wilcoxon rank-sum test was used. For the control group: *n* = 110 RGCs, *n* = 10 experiments, *n* = 9 mice. For the muscimol group, *n* = 44 RGCs, *n* = 3 experiments, *n* = 3 mice.

## REFERENCES

1. Ahissar, E., Vaadia, E., Ahissar, M., Bergman, H., Arieli, A., & Abeles, M. (1992). Dependence of Cortical Plasticity on Correlated Activity of Single Neurons and on Behavioral Context. Science, 257(5075), 1412–1415.

2. Azouz, R., & Gray, C. M. (2000). Dynamic spike threshold reveals a mechanism for synaptic coincidence detection in cortical neurons in vivo. Proceedings of the National Academy of Sciences of the United States of America, 97(14), 8110–8115.

3. Baden, T., Berens, P., Franke, K., Roson, M. R., Bethge, M., & Euler, T. (2016). The functional diversity of retinal ganglion cells in the mouse. Neture, 529, 345–350.

4. Balmer, T. S., & Pallas, S. L. (2015). Visual experience prevents dysregulation of GABAB receptor-dependent short-term depression in adult superior colliculus. Journal of Neurophysiology, 113, 2049–2061.

5. Basso, M. A., Bickford, M. E., & Cang, J. (2021). Unraveling circuits of visual perception and cognition through the superior colliculus. Neuron, 109(6), 918–937.

6. Bereshpolova, Y., Stoelzel, C. R., Gusev, A. G., Bezdudnaya, T., & Swadlow, H. A. (2006). The Impact of a Corticotectal Impulse on the Awake Superior Colliculus. The Journal of Neuroscience, 26(8), 2250–2259.

7. Bliss, T. V. P., & Gardner-Medwin, A. R. (1973). Long-lasting potentiation of synaptic transmission in the dentate area of the unanaestetized rabbit following stimulation of the perforant path. The Journal of Physiology, 232(2), 357–74.

8. Bliss, T. V. P., & Lomo, T. (1973). Long-lasting potentiation of synaptic transmission in the dentate area of the anaesthetized rabbit following stimulation of the perforant path. The Journal of Physiology, 232(2), 331–356.

9. Chance, F. S., Abbott, L. F., & Reyes, A. D. (2002). Gain modulation from background synaptic input. Neuron, 35, 773–782.

10. Chavez-Noriega, L. E., Halliwell, J. V., & Bliss, T. V. P. (1990). A decrease in firing threshold observed after induction of the EPSP-spike (E-S) component of long-term potentiation in rat hippocampal slices. Experimental Brain Research, 79, 633–641.

11. Chung, S., Li, X., & Nelson, S. B. (2002). Short-term depression at thalamocortical synapses contributes to rapid adaptation of cortical sensory responses in vivo. Neuron, 34(3), 437–446.

12. Constantinidis, C., & Goldman-Rakic, P. S. (2002). Correlated Discharges Among Putative Pyramidal Neurons and Interneurons in the Primate Prefrontal Cortex. Journal of Neurophysiology, 88(6), 3487–3497.

13. Cook, D. L., Schwindt, P. C., Grande, L. A., & Spain, W. J. (2003). Synaptic depression in the localization of sound. Nature, 421(6918), 66–70.

14. Denman, D. J., Siegle, J. H., Koch, C., Reid, R. C., & Blanche, T. J. (2017). Spatial Organization of Chromatic Pathways in the Mouse Dorsal Lateral Geniculate Nucleus. The Journal of Neuroscience, 37(5), 1102–1116.

15. English, D. F., McKenzie, S., Evans, T., Kim, K., Yoon, E., & Buzsáki, G. (2017). Pyramidal Cell-Interneuron Circuit Architecture and Dynamics in Hippocampal Networks. Neuron, 96(2), 505–520.

16. Evans, D. A., Stempel, A. V., Vale, R., Ruehle, S., Lefler, Y., & Branco, T. (2018). A synaptic threshold mechanism for computing escape decisions. Nature, 558(7711), 590–594.

17. Ferguson, K. A., & Cardin, J. A. (2020). Mechanisms underlying gain modulation in the cortex. Nature Reviews Neuroscience, 21(2), 80–92.

18. Fortune, E. S., & Rose, G. J. (2001). Short-term synaptic plasticity as a temporal filter. Trends in Neurosciences, 24(7), 381–385.

19. Gehr, C., Sibille, J., & Kremkow, J. (2023). Retinal input integration in excitatory and inhibitory neurons in the mouse superior colliculus in vivo. eLife, 12, RP88289.

20. Ghanbari, A., Ren, N., Keine, C., Stoelzel, C., Englitz, B., Swadlow, H. A., & Stevenson, I. H. (2020). Modeling the Short-Term Dynamics of in Vivo Excitatory Spike Transmission. The Journal of Neuroscience, 40(21), 4185–4202.

21. Henze, D. A., Wittner, L., & Buzsáki, G. (2002). Single granule cells reliably discharge targets in the hippocampal CA3 network in vivo. Nature Neuroscience, 5(8), 790–795.

22. Isa, T., Marquez-Legorreta, E., Grillner, S., & Scott, E. K. (2021). The tectum/superior colliculus as the vertebrate solution for spatial sensory integration and action. Current Biology, 31, R741–R762.

23. Ito, S., & Feldheim, D. A. (2018). The mouse superior colliculus: An emerging model for studying circuit formation and function. Frontiers in Neural Circuits, 12, 10.

24. Jouhanneau, J.-S., Kremkow, J., Dorm, A. L., & Poulet, J. F.A. (2015). In Vivo Monosynaptic Excitatory Transmission between Layer 2 Cortical Pyramidal Neurons. Cell Reports, 13, 2098–2106.

25. Jun, J. J., Steinmetz, N. A., Siegle, J. H., Denman, D. J., Bauza, M., Barbarits, B., Lee, A. K., Anastassiou, C. A., Andrei, A., Aydın, Ç., Barbic, M., Blanche, T. J., Bonin, V., Couto, J., Dutta, B., Gratiy, S. L., Gutnisky, D. A., Häusser, M., Karsh, B.,… Harris, T. D. (2017). Fully integrated silicon probes for high-density recording of neural activity. Nature, 551(7679), 232–236.

26. Kuba, H., Koyano, K., & Ohmori, H. (2002). Synaptic depression improves coincidence detection in the nucleus laminaris in brainstem slices of the chick embryo. European Journal of Neuroscience, 15(6), 984–990.

27. Liu, Y., Savier, E. L., DePiero, V. J., Chen, C., Schwalbe, D. C., Abraham-Fan, R.-J., Chen, H., Campbell, J. N., & Cang, J. (2023). Mapping visual functions onto molecular cell types in the mouse superior colliculus. Neuron, 111, 1876–1886.e5.

28. Marder, C. P., & Buonomano, D. V. (2003). Differential Effects of Short-and Long-Term Potentiation on Cell Firing in the CA1 Region of the Hippocampus. The Journal of Neuroscience, 23(1), 112–121.

29. Martinetti, L. E., Bonekamp, K. E., Autio, D. M., Kim, H.-H., & Crandall, S. R. (2022). Short-Term Facilitation of Long-Range Corticocortical Synapses Revealed by Selective Optical Stimulation. Cerebral Cortex, 32(9), 1932–1949.

30. Mongillo, G., Barack, O., & Tsodyks, M. (2008). Synaptic theory of working memory. Science, 319(5869), 1543–1546.

31. Pachitariu, M., Steinmetz, N., Kadir, S., Carandini, M., & Harris, K. D. (2016). Kilosort: realtime spike-sorting for extracellular electrophysiology with hundreds of channels. bioRxiv, 061481.

32. Pierce, J. W. (2008). Generating stimuli for neuroscience using PsychoPy. Frontiers in Neuroinformatics, 2, 10.

33. Sauvé, Y., Sawai, H., & Rasminsky, M. (1995). Functional synaptic connections made by regenerated retinal ganglion cell axons in the superior colliculus of adult hamsters. The Journal of Neuroscience, 15(1), 665–675.

34. Shamash, P., Carandini, M., Harris, K. D., & Steinmetz, N. (2018). A tool for analyzing electrode tracks from slice histology. bioRxiv, 447995.

35. Shein-Idelson, M., Pammer, L., Hemberger, M., & Laurent, G. (2017). Large-scale mapping of cortical synaptic projections with extracellular electrode arrays. Nature Methods, 14, 882–890.

36. Shi, X., Barchini, J., Ledesma, H. A., Koren, D., Jin, Y., Liu, X., Wei, W., & Cang, J. (2017). Retinal origin of direction selectivity in the superior colliculus. Nature Neuroscience, 20(4), 550–558.

37. Sibille, J., Gehr, C., Benichov, J. I., Balasubramanian, H., Teh, K. L., Lupashina, T., Vallentin, D., & Kremkow, J. (2022a). High-density electrode recordings reveal strong and specific connections between retinal ganglion cells and midbrain neurons. Nature Communications, 13, 5218.

38. Sibille, J., Gehr, C., & Kremkow, J. (In press). Efficient mapping of the thalamocortical monosynaptic connectivity in vivo by tangential insertions of high-density electrodes in the cortex. Proceedings of the National Academy of Sciences.

39. Sibille, J., Gehr, C., Teh, K. L., & Kremkow, J. (2022b). Tangential high-density electrode insertions allow to simultaneously measure neuronal activity across an extended region of the visual field in mouse superior colliculus. Journal of Neuroscience Methods, 376, 109622.

40. Stark, E., & Abeles, M. (2009). Unbiased estimation of precise temporal correlations between spike trains. Journal of Neuroscience Methods, 179(1), 90–100.

41. Steinmetz, N. A., Zatka-Haas, P., Carandini, M., & Harris, K. D. (2019). Distributed coding of choice, action and engagement across the mouse brain. Nature, 576, 266–273.

42. Stevenson, I. H. (2023). Circumstantial evidence and explanatory models for synapses in large-scale spike recordings. *Neurons, Behavior*, Data Analysis, and Theory, 1–30.

43. Stoelzel, C. R., Bereshpolova, Y., Gusev, A. G., & Swadlow, H. A. (2008). The Impact of an LGNd Impulse on the Awake Visual Cortex: Synaptic Dynamics and the Sustained/Transient Distinction. The Journal of Neuroscience, 28(19), 5018–5028.

44. Taube, J. S., & Schwartzkroin, P. A. (1988). Mechanisms of long-term potentiation: EPSP/spike dissociation, intradendritic recordings, and glutamate sensitivity. The Journal of Neuroscience, 8(5), 1632–1644.

45. Tominaga, T., & Tominaga, Y. (2016). Paired Burst Stimulation Causes GABAA Receptor-Dependent Spike Firing Facilitation in CA1 of Rat Hippocampal Slices. Frontiers in Cellular Neuroscience, 10, 9.

46. Usrey, W. M., Reppas, J. B., & Reid, R. C. (1998). Paired-spike interactions and synaptic efficacy of retinal inputs to the thalamus. Nature, 395, 384–387.

47. Veinante, P., & Deschênes, M. (2003). Single-cell study of motor cortex projections to the barrel field in rats. The Journal of Comparative Neurology, 464(1), 98–103.

